# Essentially leading antibody production: An investigation of amino acids, myeloma and natural V-region signal peptides in producing Pertuzumab and Trastuzumab variants

**DOI:** 10.1101/2020.07.27.221887

**Authors:** Wei-Li Ling, Chinh Tran-To Su, Wai-Heng Lua, Jun-Jie Poh, Yuen-Ling Ng, Anil Wipat, Samuel Ken-En Gan

## Abstract

Boosting the production of recombinant therapeutic antibodies is crucial in both academic and industry settings. In this work, we investigated the usage of varying signal peptides by antibody genes and their roles in recombinant transient production. Comparing myeloma and the native signal peptides of both heavy and light chains in 168 antibody permutation variants, we performed a systematic analysis, finding amino acids counts to be involved in antibody production to construct a model for predicting co-transfection transient recombinant antibody production rates using the HEK293 system. The findings also provide insights into the usage of the large repertoire of antibody signal peptides.

## 1 Introduction

The signal peptide (SP) of a protein is a short tag of amino acids at the N- or C-terminal that predestinates the protein location extracellularly or within the cell to the organelles. Known organelle targeting SPs include the nucleus localization (Kalderon et al., 1984) or export signal (Wen et al., 1995), mitochondria signals (Maccecchini et al., 1979), endoplasmic reticulum (ER) secretion (Blobel and Dobberstein, 1975) or retention signal (Stornaiuolo et al., 2003)), and peroxisome signals (Gould et al., 1987).

Secreted by plasma B cells, antibodies are tagged with the ER secretion signal/SP at the N-terminal and translocated into the ER lumen (Walter et al., 1981), before being passed to the Golgi apparatus and sorted into secretory vesicles for extracellular secretion (Beams and Kessel, 1968). The SP is unique to the protein and generally contains a positively charged N-terminal, followed by a hydrophobic region and a neutral polar C-terminal (von Heijne and Gavel, 1988). Depending on its location at the N or C terminal of the protein, there is a cleavage site separating the SP from the protein that would be recognized by a signal peptidase (Milstein et al., 1972). While secretory SPs are involved in the co-translational co-translocation pathway (Blobel and Dobberstein, 1975) with the primary function to export proteins (Kapp et al., 2000), it remains enigmatic to why antibodies utilize a large repertoire of SPs (e.g. Vκ1 family with 22 SPs, VH3 family with 50 SPs, retrieved from the IMGT database (Giudicelli et al., 2005) at the point of writing), hinting of possible roles in antibody production.

Modifying SPs for better recombinant protein production has been largely successful in many studies (Haryadi et al., 2015; Huang et al., 2017; Ohmuro-Matsuyama and Yamaji, 2018; Zhou et al., 2016), however, the underlying mechanism of such effects remains enigmatic. With recent studies demonstrating cross-talks of antibody elements, where the constant regions (Lua et al., 2018; Su et al., 2018), variable regions and their pairings (Ling et al., 2018; Lua et al., 2019) affect antibody production and function, there is increasing evidence that antibodies ought to be investigated holistically (Phua et al., 2019), especially as therapeutics (Ling et al., 2020). Through the inclusion of antibody SP in the analysis, deeper insights into antibody V-region pairing effects in transient recombinant production (Ling et al., 2018) can be further holistically considered. In this work, we unraveled the role of total amino acid usage that may underlie the usage of the diverse repertoire of antibody SPs in compensating for V-region hypervariability to overcome production bottlenecks.

## 2 Materials and Methods

### 2.1 Signal peptide selection

SP data were retrieved from the IMGT database for the respective light and heavy chain families. Consensus sequences within each family SP were determined using WebLogo (Crooks et al., 2004) (https://weblogo.berkeley.edu/logo.cgi) to derive the VH and Vκ family SPs.

### 2.2 Recombinant antibody production

All VH and Vκ sequences used were described previously (Ling et al., 2018) and codon optimized using GenSmart™ Codon Optimization (Genscript) online software, followed by synthesis by Eurofin (Japan) to avoid codon usage bias. The genes were transformed into competent E. coli (DH5α) strains (Chan et al., 2013) followed by plasmid extraction (Biobasic Pte Ltd) and sub-cloning into pTT5 vector (Youbio) using restriction enzyme sites, as previously performed (Ling et al., 2018; Lua et al., 2018; Lua et al., 2019; Su et al., 2017).

### 2.3 Site Directed mutagenesis and SP swapping

Mutations in Vκ1 SP were performed using QuikChange Lightning Site-Directed Mutagenesis Kit (Agilent) with the respective primers: P18S forward: 5’- CTG CTG CTG CTT TGG CTT TCT GGC GCT AG-3’; P18S reverse: 5’-CTA GCG CCA GAA AGC CAA AGC AGC AGC AG-3’; P18R forward: 5’- GCT GCT TTG GCT TCG TGG CGC TAG ATG CG-3’; P18R reverse: 5’- CGC ATC TAG CGC CAC GAA GCC AAA GCA GC-3’ and protocols according to manufacturer recommendations.

SP graftings were performed using overhanging primers to the various V-genes via PCR using Q5 polymerase (NEB) with the following primers: VH3 18P 1st leader extension forward: 5’-CAG CTG CTG GGC CTG CTG CTG CTT TGG CTT CCT GGC GCT AGA TGC GAA GTG CAG CTG GTG-3’; VH3 18R 1st leader extension forward: 5’-CAG CTG CTG GGC CTG CTG CTG CTT TGG CTT CGT GGC GCT AGA TGC GAA GTG CAG CTG GTG-3’; VH3 18S 1st leader extension forward: 5’ - CAG CTG CTG GGC CTG CTG CTG CTT TGG CTT TCT GGC GCT AGA TGC GAA GTG CAG CTG GTG-3’; VH3 18P 2nd leader extension forward: 5’ - GCC GAA TTC GCG GCC GCG TTC CTC ACC ATG GAC ATG AGA GTT CCA GCT CAG CTG CTG GGC-3’; VH3 leader extension reverse: 5’-GCC GCG AAG CTT TCA CTT GCC AGG AGA-3’.

### 2.4 Bio-Layer Interferometry quantification

The Octet Red96 system (ForteBio) was used to quantify the amount of antibodies in transiently co-transfected cell cultures supernatants using Protein G biosensors (ForteBio) with preloaded program settings (high sensitivity assay with regeneration) in Octet Data Acquisition v10.0 as previously described (Ling et al., 2018; Lua et al., 2018; Lua et al., 2019; Su et al., 2017).

Quantification data were analyzed using Octet data analysis v10.0 with protein standard ranged from 100 μg/ml to 0.1953 μg/ul in two-fold serial dilution as per described in (Ling et al., 2018; Su et al., 2017).

### 2.5 Constructing the statistical predictive model

The logistic regression model was built from the data from present and previous works (Ling et al., 2018). Production rates were categorized into low production (< 20%), medium (from 20% to 70%) and high (> 70%) normalized by datasets of the IgE SP. The Logistic Regression CV classifier using “lbfgs” optimizer with L2 norm (implemented in the scikit learn v.0.22.1 package (Pedregosa et al., 2011)) was used in a 20-fold cross validation process to fine-tune the regularization strength. A weighted precision scoring function was used to evaluate the model performance and the categorized classes (low, medium, high) were weighted to counter the slight imbalance in the dataset, i.e. 65% low, 70% medium, and 33% high production samples. The optimization process was performed in 1000 iterations.

The probability of each categorized production label (class label) of each antibody variant was calculated using the equation below:

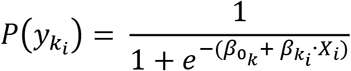

where:

*p*(*y_k_i__*) is probability of the predicted class label yk (i.e. the production level) of variant i, with k = 0 (low), 1 (medium), 2 (high)
*β*_0_*k*__ is the corresponding intercept of class k, with β0 = [−17.88, 14.62, 3.25] with respect to k = [0, 1, 2]
*β_k_i__* is array of regression coefficients of features (amino acid contribution) in variant i for each class k
*X_i_* is array of amino acid counts in each variant i

To evaluate the model performance, average area under ROC (AUC) of all pairwise combinations of classes and averaged F1-score were computed. In addition, a dummy model using Dummy classifier with default parameters was created and used as a baseline control.

## 3 Results

### 3.1 “IgE signal peptide” results in better antibody production rates

Antibody SPs were initially named by the constant region isotypes but were recently re-classified in IMGT by the V-region family. With exception to the wild-type IgE signal peptide: Humighae 1 (Genbank accession J00227), termed ‘IgE’ SP, references to the SPs in this research follows the IMGT convention. Given the large number of antibody germline SPs, we selected consensus/dominant representatives of each VH and Vκ family as representative ‘native’ SPs.

VH and Vκ variants with Pertuzumab and Trastuzumab CDRs were paired with their respective SPs (i.e. VH1 SP to VH1 framework (FWR), Vκ1 SP to Vκ1 FWR) and production levels compared to utilizing only the IgE SP from our previous work (Ling et al., 2018). Given the variability in transient co-transfections, recombinant Pertuzumab and Trastuzumab Vκ1 VH3 with the IgE SP were used for normalization (100%) to facilitate comparisons. Given the high CDR similarity of both Trastuzumab and Pertuzumab (Ling et al., 2018), this allows an investigation of minute CDR difference effects on protein production.

Within the Pertuzumab variants (Figures 1A & C), production levels with the IgE SP (Ling et al., 2018) tend to be higher than those using native SPs (e.g. highest producing pair using the IgE SP: Vκ4|VH3 at 125%, while the highest producing pair utilizing their respective SP, were Vκ3|VH7 at 80%) with exceptions (Vκ1|VH1, Vκ2|VH1, Vκ1|VH4, Vκ2|VH4, Vκ1|VH7, and Vκ2|VH7). Regardless of the SP, antibodies paired with Vκ5 family constructs gave poor yields (Figure 1C, sorted by light chain families).

**Figure 1.**
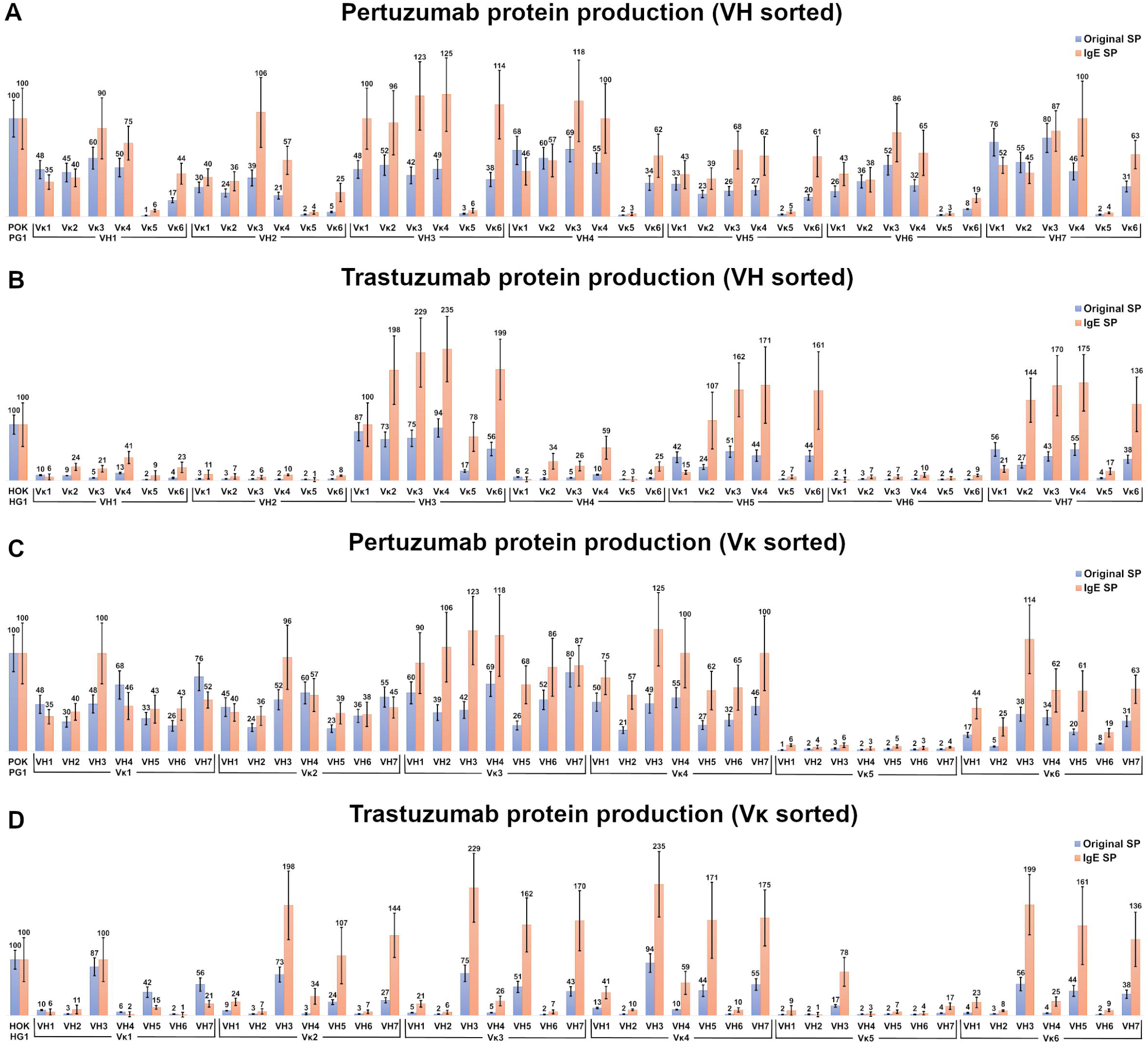
Production rates (%) of recombinant antibodies containing various heavy and light chain pairings. (A & B) Recombinant Pertuzumab (A) and Trastuzumab (B) variants with the various SPs grouped according to VH families. (C & D) Recombinant Pertuzumab and Trastuzumab variants with the various SPs grouped according to Vκ families. Recombinant Pertuzumab and Trastuzumab variants produced with native (blue) or the ‘IgE’ (orange) SPs respectively, are shown. In all experiments, the recombinant wild-type Pertuzumab (POK PG1) or Trastuzumab (HOK HG1) was used (shown in the first column). The production rate of the variants utilizing the ‘IgE’ SP (Ling et al., 2018) was used as the reference for the production rate.

Of the Trastuzumab variants (Figure 1B & D), there is general agreement to the trends observed in the Pertuzumab dataset where IgE SP variants generally having higher production rates than native SP variants. The highest producing antibody with IgE SP is Vκ4|VH3 at 235%, with its corresponding counterparts with native SPs at 94%. The only exceptions where the native SPs had higher productions were the Vκ1 |VH5 and Vκ1 |VH7 pairs.

In the Pertuzumab dataset, the Vκ5 family is the sole poor producing family, whereas the low production families in the recombinant Trastuzumab model dataset (Figure 1D) extended to VH1, 2, 4 and 6 (light chains that paired with these VHs had lower yields). One notable exception was the Vκ5|VH3 pair that was produced at higher levels compared to other Vκ5 family permutations in the Trastuzumab dataset (Figure 1D).

VH1 and VH7 genes shared the same SP (as classified in IMGT). amino acid sequence but with different codons. While this was normalized through codon optimization, there were still distinct different productions between the two VH families, with VH7 being the better producing partner (average production of VH7 with its light chain partners at 48% when using native SPs and at 58% when using IgE SP) compared to VH1 (at 37% when using native SPs and at 48% when using IgE SP) in the Pertuzumab dataset. The effect between VH1 and VH7 was even more pronounced in the Trastuzumab dataset, where VH1 recombinant production level were significantly reduced (at 7% when using native SPs, and 20% when using IgE SP) as compared to VH7 (at 37% when using native SPs, and 110% when using the IgE SP).

### 3.2 Comparison of IgE, Vκ1 and native SPs in recombinant Pertuzumab and Trastuzumab antibody production

To investigate the role of light chain SPs, Vκ1 SP was grafted onto the Pertuzumab and Trastuzumab VH3 FWR and compared to the IgE and native SPs pairings after normalization with the respective Trastuzumab/Pertuzumab variants with the IgE SP (Figure 2). The respective heavy and light chains of Pertuzumab/Trastuzumab with the IgE SP are termed IgESP-Vκ1 (light chain) and IgESP-VH3 (heavy chain) in Figure 2. The native SPs (Vκ1SP-Vκ1, VH3SP-VH3) had the lowest recombinant production at 48% and 87% for the Pertuzumab and Trastuzumab models, respectively. The Vκ1 SP (Vκ1SP-Vκ1, Vκ1SP-VH3) had the highest production at 112% and 240% in the Pertuzumab and Trastuzumab models, respectively.

**Figure 2.**
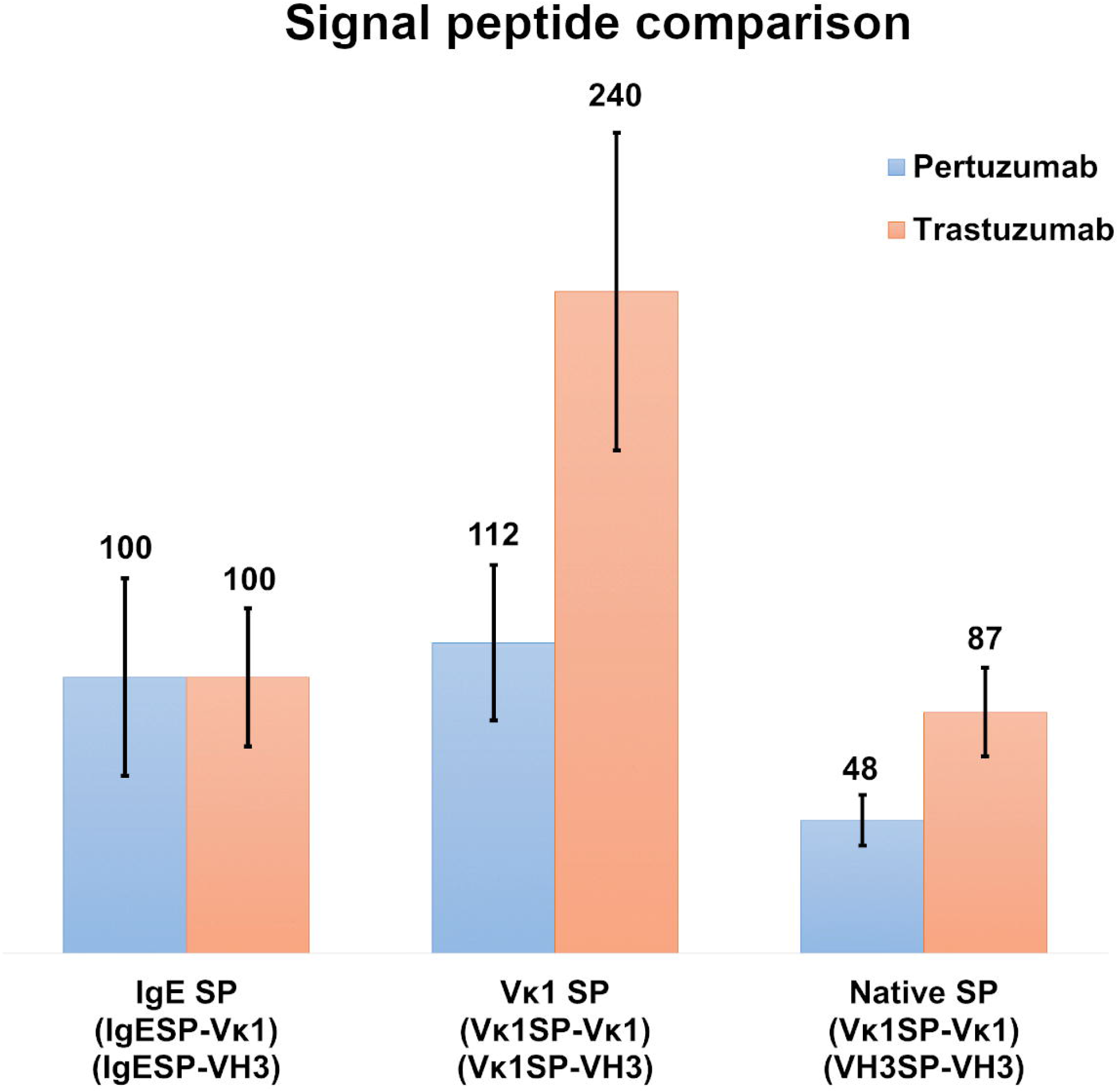
Transient recombinant antibody production rates of wild-type Vκ1|VH3 Pertuzumab (blue) and Trastuzumab (orange) using the IgE, Vκ1 and native SPs in %.

### 3.3 Myeloma SPs production rates

To study the effect of myeloma SPs on improving production levels, we performed single amino acid mutagenesis (P18R and P18S) on the Vκ1 SP. The mutated SPs were generated based on a previous reported myeloma Vκ1 SP associated Fanconi’s syndrome SP (Rocca et al., 1995), and the IgE SP sequence, also from a myeloma patient (Nilsson et al., 1970). The Vκ1 P18R and P18S SPs showed 56% & 55% in Pertuzumab and 70% & 170% in Trastuzumab, respectively, compared to native Vκ1 SP at 112% & 240% in Pertuzumab and Trastuzumab, respectively. Both mutations had lower productions compared to the native Vκ1 SP, but the P18S SP Trastuzumab had notably higher recombinant production than the wild-type IgE SP.

### 3.4 The roles of essential and non-essential amino acid on antibody production

With no clear correlation between production levels and the SP used, the content of the SPs was investigated. EAAs were found to make up half of the SP length (Table 1) with general varied usage, with some being more prominent, e.g. leucine (L).

**Table 1.** List of SP sequences with essential amino acids (EAAs, various colors) and non-essential amino acids (NEAAs, uncoloured) usage shown. Comparisons of the total variety counts of EAAs in the SPs explored in this study.

| Family | Signal peptide amino acid position |  |  |  |  |  |  |  |  |  |  |  |  |  |  |  |  |  |  |  |  |  | Total variety/<br>No. of<br>EAA |
| --- | --- | --- | --- | --- | --- | --- | --- | --- | --- | --- | --- | --- | --- | --- | --- | --- | --- | --- | --- | --- | --- | --- | --- |
|  | 1 | 2 | 3 | 4 | 5 | 6 | 7 | 8 | 9 | 10 | 11 | 12 | 13 | 14 | 15 | 16 | 17 | 18 | 19 | 20 | 21 | 22 |  |
| Vκ1 | M | D | M | R | V | P | A | Q | L | L | G | L | L | L | L | W | L | P | G | A | R | C | 4 / 11 |
| Vκ2 | M | R | L | P | A | Q | L | L | G | L | L | M | L | W | V | P | G | S | S | A |  |  | 4 / 10 |
| Vκ3 | M | E | A | P | A | Q | L | L | F | L | L | L | L | W | L | P | D | T | T | G |  |  | 5 / 12 |
| Vκ4 | M | V | L | Q | T | Q | V | F | I | S | L | L | L | W | I | S | G | A | Y | G |  |  | 7 / 12 |
| Vκ5 | M | G | S | Q | V | H | L | L | S | F | L | L | L | W | I | S | D | T | R | A |  |  | 8 / 12 |
| Vκ6 | M | L | P | S | Q | L | I | G | F | L | L | L | W | V | P | A | S | R | G |  |  |  | 6 / 10 |
| VH1 | M | D | W | T | W | R | I | L | F | L | V | A | A | A | T | G | A | H | S |  |  |  | 8 / 11 |
| VH2 | M | D | I | L | C | S | T | L | L | L | L | T | V | P | S | W | V | L | S |  |  |  | 6 / 13 |
| VH3 | M | E | F | G | L | S | W | V | F | L | V | A | I | L | K | G | V | Q | C |  |  |  | 7 / 12 |
| VH4 | M | K | H | L | W | F | F | L | L | L | V | A | A | P | R | W | V | L | S |  |  |  | 7 / 14 |
| VH5 | M | G | S | T | A | I | L | A | L | L | L | A | V | L | Q | G | V | C | S |  |  |  | 5 / 10 |
| VH6 | M | S | V | S | F | L | I | F | L | P | V | L | G | L | P | W | G | V | L | S |  |  | 6 / 13 |
| VH7 | M | D | W | T | W | R | I | L | F | L | V | A | A | A | T | G | A | H | S |  |  |  | 8 / 11 |
| IgE | M | D | W | T | W | I | L | F | L | V | A | A | A | T | R | V | H | S |  |  |  |  | 8 / 12 |

Light chain SPs tend to be longer in length, more homogenous (4-8 EAA) with lower EAA counts (10-12 amino acids) than the heavy chain SPs. This involvement of EAAs may elucidate the observations that the Vκ1 SP yielded better production rates than those of the IgE SP (Figures 1 & 2), which in turn, yielded better production than the native SPs.

Analysis of EAA content (Table 1) demonstrated that native SPs consisting of higher EAA counts result in lower production levels compared to IgE or Vκ1 SPs (Figure 1 & 2).

Based on the SP usage of EAA and NEAA, the usage of these amino acids may extend beyond just the SP to the whole protein. Applying this analysis to the full-length Pertuzumab and Trastuzumab variants (including light and heavy chains), we found higher counts of phenylalanine (F), histidine (H), isoleucine (I), alanine (A), asparagine, (N) and lower counts of leucine (L) and serine (S) within the Pertuzumab variants (Figure 4) which may contribute to the poor production levels of the Vκ5 family. We observed that higher counts of L, arginine (R) and lower counts of I, lysine (K), and aspartic acid (D) may account for the increased production in the Vκ3 family. Higher counts of tryptophan (W) may also account for higher production in general.

Analysis of the Trastuzumab variants (Figure 5) showed similar trends in amino acids usage due to the high similarities of CDRs between Trastuzumab and Pertuzumab. This extended to the poor production in Vκ5 of both Trastuzumab (Figure 5) and Pertuzumab (Figure 4). Higher counts of D appear to improve production in Trastuzumab Vκ4 variants, contrary to Pertuzumab Vκ3 variants. Higher counts of Tyrosine (Y), Proline (P), Glycine (G) and lower counts of Glutamine (Q) are associated with better production in our Trastuzumab repertoire.

### 3.5 The effects of amino acid supply on recombinant antibody production

Since the culture media is the predominant nutrient source in transient recombinant protein production, we analyzed the amino acid constituents and the demand for the respective amino acids. For simpler analysis, we deemed batch variations involving co-transfection procedure variations and serum differences to be negligible. Based on the DMEM formulation (Sigma Aldrich, Cat no. D1152), W was found to be the limiting amino acid (0.47 x 1020 molecules) compared to other EAAs, restricting our maximum production to 1.88 x 1018 of antibodies (a representative average of amino acid usage from all the antibodies in this study, see Supplementary Table 1-2).

Five NEAAs: A, D, E, N and P were not provided for in DMEM probably assumed to be sufficient from internal cell synthesis. However, three NEAAs: S, D, and Glutamic acid (E) are precursors to Cysteine (C), G, N, P, Q, and R but were also absent in the media, making these amino acids potential limiting factors.

### 3.6 A statistical predictive model of antibody production rate

To demonstrate the noticeable involvement of varying SP and the amino acid counts, we constructed a statistical model to computationally predict antibody production rates based on our co-transfection transient HEK293 cell system. The logistic regression model is based on the data in this study and includes other antibodies from our previous work (Ling et al., 2018) for better machine learning.

The prediction scores of AUC ~0.79–0.95 (Figure 6) and F1-score 0.62–0.7 (of which 1 reflecting the best balance between precision and recall) depicted a reasonable prediction model of production rate categories (low, medium, or high) using various signal peptide sequences.

## 4 Discussion

We investigated the effects of the numerous antibody signal peptides (SP) on recombinant antibody production in a co-transfection transient system using HEK293 cells, normalizing with the same transfection agents and backbone plasmids to varying only the signal peptides. We found that the IgE SP generally gave better yields than the native SPs (Figure 1).

To study possible effects of SPs from heavy and light chains (Haryadi et al., 2015), we grafted the Vκ1 SP on the wild-type Vκ1|VH3 of the recombinant Pertuzumab and Trastuzumab (Figure 2) for comparison to IgE SP and the respective native SP counterparts. Both IgE and Vκ1 SPs yielded better productions. With the possibility that myeloma SP might give better production, we studied another myeloma-linked SP – a variant of Vκ1 SP -with P18R or P18S different from the Vκ1 SP in IMGT (Rocca et al., 1995).

Grafting of the myeloma Vκ1 SP onto the wild-type Vκ1 |VH3 recombinant Pertuzumab and Trastuzumab models showed that mutation P18S increased production only in the Trastuzumab model, while the other mutation had significantly lower production than the native Vκ1 SP (Figure 3). Multiple sequence alignment analysis (Supplementary Figure 1 & 2) of the various native SPs showed major similarity in amino acid usage and positions across the SPs of different families. The hydrophobic center region of the SPs was shown to be conserved, while the N- and C-terminal regions were more distinctive. Given these findings, we did not find support for the role of SP in hyperglobulinemia pathogenesis within myeloma, and that there remains an enigmatic relationship between the SP and antibody production.

**Figure 3.**
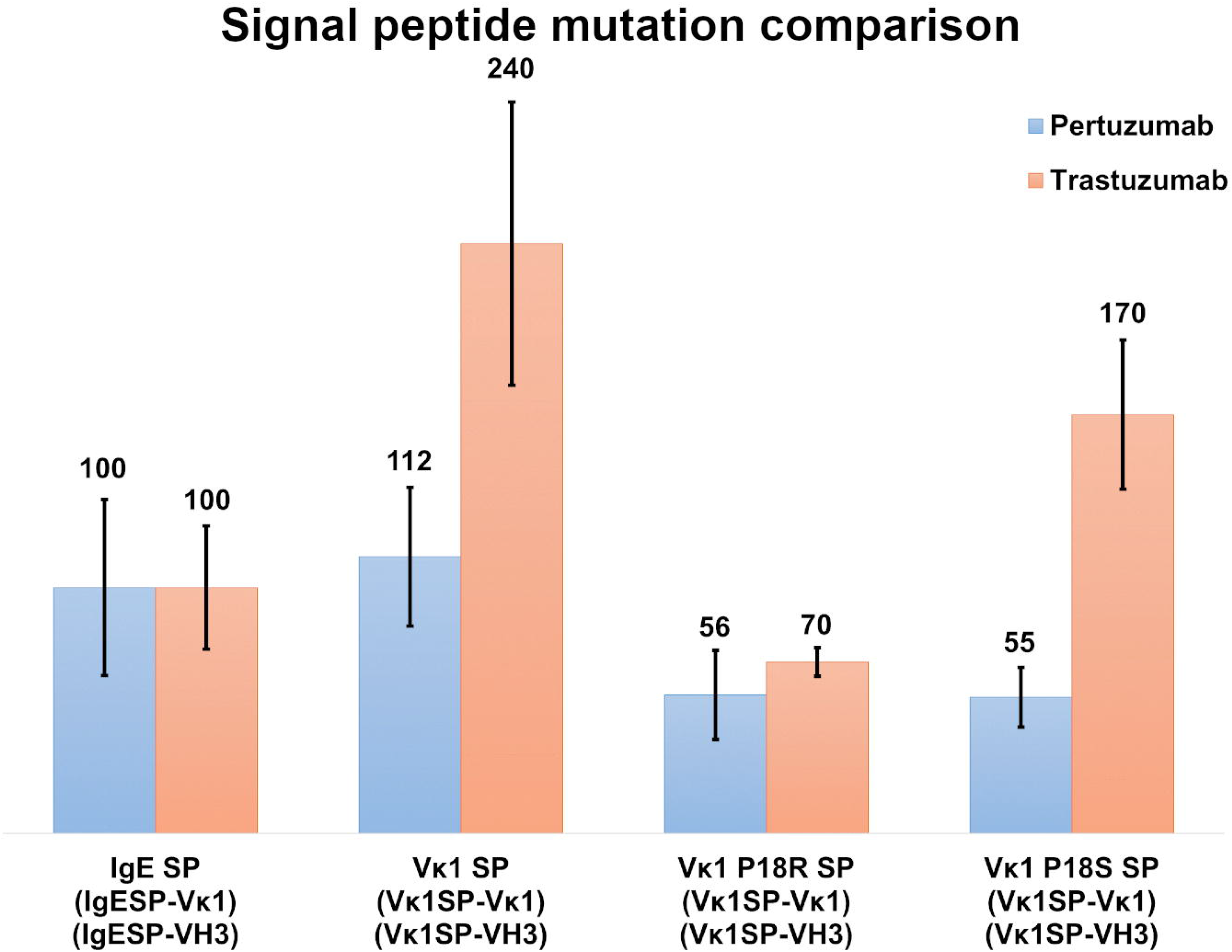
Recombinant protein production (%) with IgE, Vκ1 and mutated Vκ1 SPs, (Vκ1 P18R and Vκ1 P18S) against the wild-type Vκ1|VH3 Pertuzumab (blue) and Trastuzumab (orange) models.

Analyzing the nine essential amino acid (EAA) content in SP sequences, especially given that previous experiments showed that the addition of supplements may be helpful in protein production, we found that light chain SPs had a lower variety and number of EAAs than the heavy chain SPs (Table 1), providing a clue to their possible contribution to recombinant production.

Extending the essential and non-essential amino acid analysis (Figure 4 and 5) to both full-length Pertuzumab and Trastuzumab variants, we found that the presence of F, H, I, A, N, L and S amino acids may underlie the poor production of Vκ5 Pertuzumab and Trastuzumab variants. On the other hand, L, R, I, K and D amino acids may allow for better production as seen with specific variants paired with Vκ3 (Pertuzumab) and Vκ4 (Trastuzumab).

**Figure 4.**
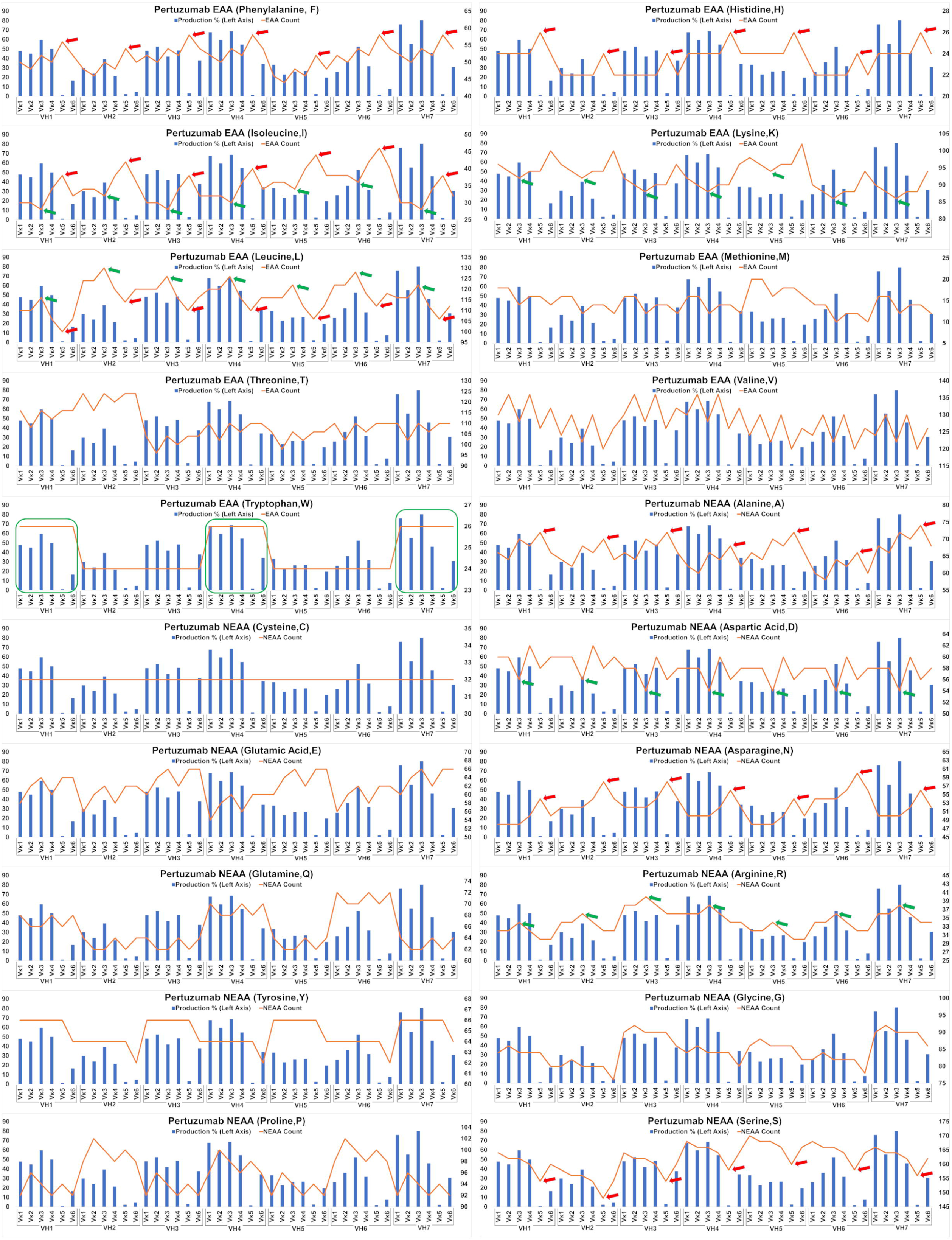
Combinations of recombinant transient recombinant antibody production in % (in bars, left axis) and amino acid counts (line, right axis) charts for the comparison to amino acids counts in recombinant production of Pertuzumab variants. The green arrows/boxes refer to an increase in production level while the red arrows depict a decrease in production levels.

**Figure 5.**
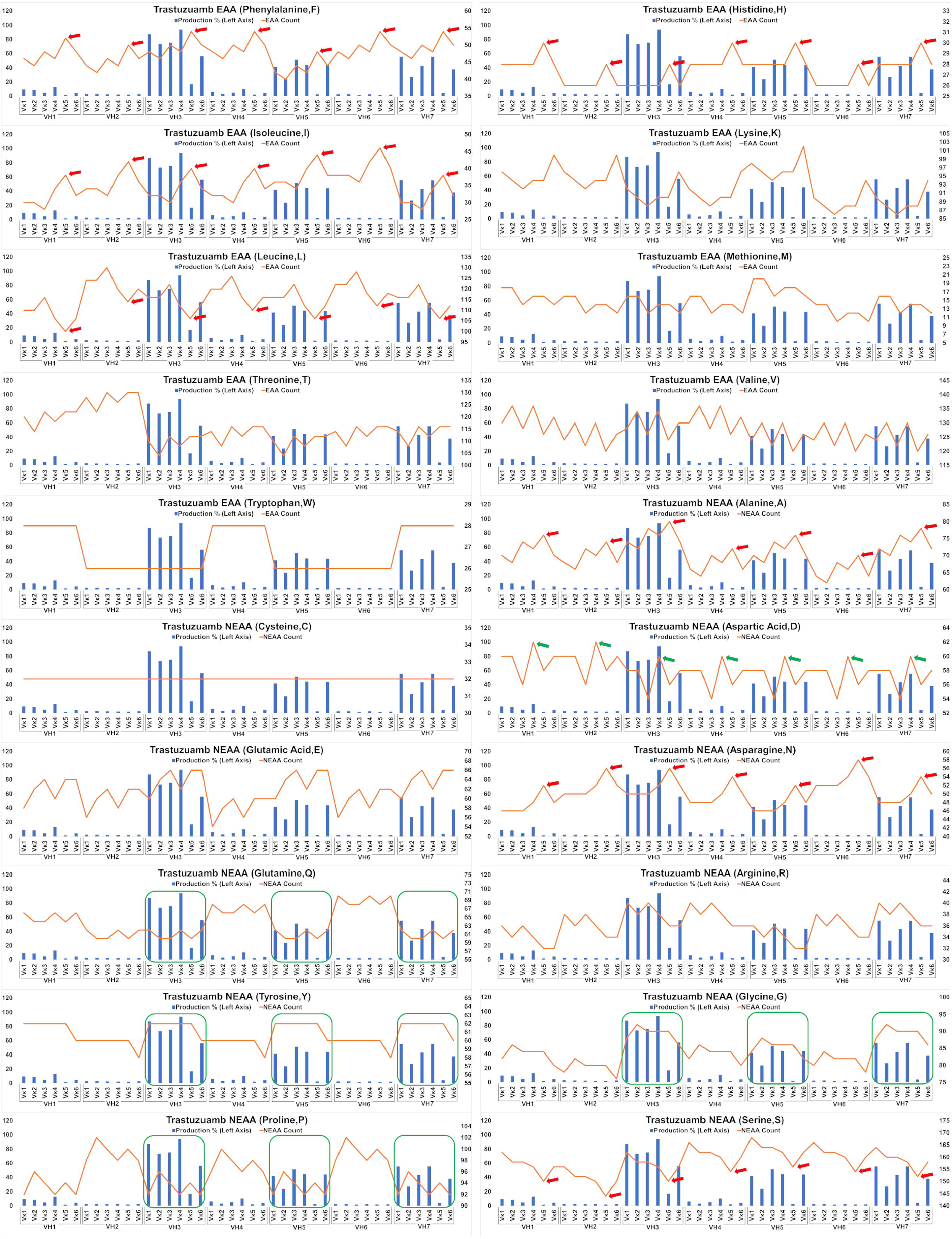
Combination of recombinant transient recombinant antibody productions in % (in bars, left axis) and amino acid counts (line, right axis) charts for the comparison to amino acids in recombinant production in Trastuzumab variants. The green arrows/boxes refer to an increase in production level while the red arrows depict a decrease in production levels.

Considering DMEM formulation (Table 2) on assumptions of negligible differences between batches of serum and co-transfected cell batches, we found that DMEM media had insufficient amino acids to support optimal antibodies production, especially since EEA: W, and NEAAs: A, D, E, N, and P, were not provided at all nor present in inadequate quantities. This thus explains the boost in production when supplements such as peptone and casein are added (Davami et al., 2014; Heidemann et al., 2000; Michiels et al., 2011; Pham et al., 2005).

**Table 2.** Amino acid counts in DMEM media and the representative average of amino acid usage in all the antibodies in this study. The number of amino acids in DMEM media were calculated based on the weight (g/L) of amino acid component used in the formula. Maximum number of antibodies produced refer to the theoretical maximum number of full-length antibodies that could be synthesized based on number of amino acids in DMEM media and to the amino acid counts in the representative average of amino acids.

| | | AA counts in full-length antibody | Number of AA in DMEM media ($10^{20}$) | Maximum number of possible antibodies produced ($10^{18}$) |
| --- | --- | --- | --- | --- |
| EAAs | Phenylalanine (F) | 50 | 2.41 | 4.82 |
|  | Histidine (H) | 26 | 1.63 | 6.27 |
|  | Isoleucine (I) | 34 | 4.82 | 14.18 |
|  | Lysine (K) | 92 | 6.01 | 6.53 |
|  | Leucine (L) | 116 | 4.82 | 4.16 |
|  | Methionine (M) | 14 | 1.21 | 8.64 |
|  | Threonine (T) | 112 | 4.80 | 4.29 |
|  | Valine (V) | 128 | 4.83 | 3.77 |
|  | Tryptophan (W) | 25 | 0.47 | 1.88 |
| NEAAs | Alanine (A) | 68 | 0 | 0 |
|  | Tyrosine (Y) | 62 | 3.45 | 5.56 |
|  | Serine (S) | 160 | 2.41 | 1.51 |
|  | Cysteine (C) | 32 | 3.11 | 9.72 |
|  | Glycine (G) | 84 | 2.41 | 2.87 |
|  | Aspartic Acid (D) | 58 | 0 | 0 |
|  | Asparagine (N) | 51 | 0 | 0 |
|  | Glutamic Acid (E) | 62 | 0 | 0 |
|  | Proline (P) | 96 | 0 | 0 |
|  | Glutamine(Q) | 63 | 24.07 | 38.21 |
|  | Arginine (R) | 35 | 2.91 | 8.31 |

Our predictive model based on amino acid accounts (Figure 6) was able to predict the ordinal production levels (low, medium, or high) with high accuracy (ROC AUC ~0.79–0.95). Nonetheless, the model is sensitive to the imbalanced datasets (e.g. 15-20% of the ‘high’ producers) despite the use of a few weighted parameters. In addition, it is confined by the transient expression conditions used in our experiments. Nonetheless, other similar transfection data can be incorporated to further improve the model for wider applicability in future work.

**Figure 6.**
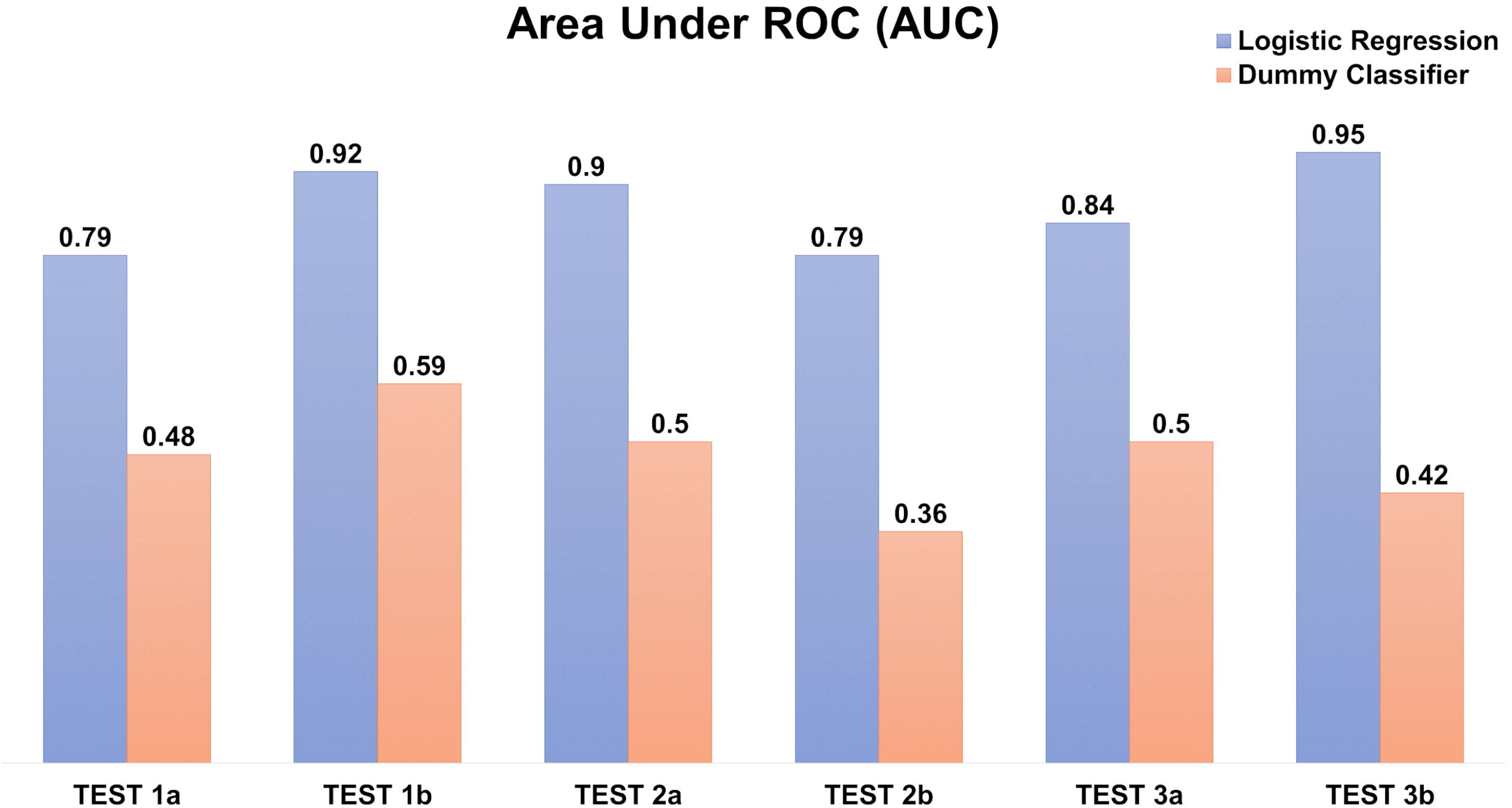
Logistic regression model to predict the recombinant antibody production rate using amino acid counts. The prediction is evaluated using Area under the ROC (AUC) in triplicates, each with two independents (a, b) testing datasets. The ‘dummy’ classifier is used as a baseline control.

With amino acid counts able to predict antibody production rates, the underlying reason that antibody genes utilize such a big repertoire of SPs may be rationalized. It is possible that the repertoire may serve to mitigate over reliance of specific EAAs that impact antibody production. Considering that the hypervariable antibody VDJ genes (Tonegawa, 1983) utilize a wide permutation of the amino acids, a fixed signal peptide for all antibodies can increase the probability of heavy bias towards specific amino acids. Such biases can thus create bottlenecks to certain EAA(s) and hamper not only antibody production, but also essential cellular protein production. Therefore, by varying the SPs with differing EAA content, such bottlenecks of EAAs may be mitigated, thus explaining a function of the large repertoire of SPs in antibody genes.

In conclusion, the study of antibodies SP with EAA factors provides a new approach on the understanding of transient mammalian protein production and possible insights to the repertoire of SPs utilized by antibodies. These new EAA factors could have potential for wide-spread application to other production systems in a better understanding of protein production.

## Conflict of Interest

The authors declare that the research was conducted in the absence of any commercial or financial relationships that could be construed as a potential conflict of interest.

## Author Contributions

Conceptualization, W.L.L. and S.K.G.; Methodology, W.L.L.; Investigation, W.L.L. and C.T.T.S.; Validation, W.H.L. and J.J.P.; Writing – Original Draft, W.L.L. and C.T.T.S.; Writing – Review & Editing, W.L.L., C.T.T.S., Y.L.N. and S.K.G.; Funding Acquisition, S.K.G.; Supervision, S.K.G., Y.L.N. and A.W.

## Funding

This work was supported by the Interstellar Initiative 2018-2019 jointly supported by the Japan Agency for Medical Research and Development and New York Academy of Sciences, and also EDDC, A*STAR.

## Notes

### Competing Interest Statement

The authors have declared no competing interest.

### Summary of Updates

Title and format revision

