## Supplementary File for "Essentially leading antibody production: An investigation of amino acids, myeloma and natural V-region signal peptides in producing Pertuzumab and Trastuzumab variants"

### *Supplementary Material*

#### 1 Supplementary Figures

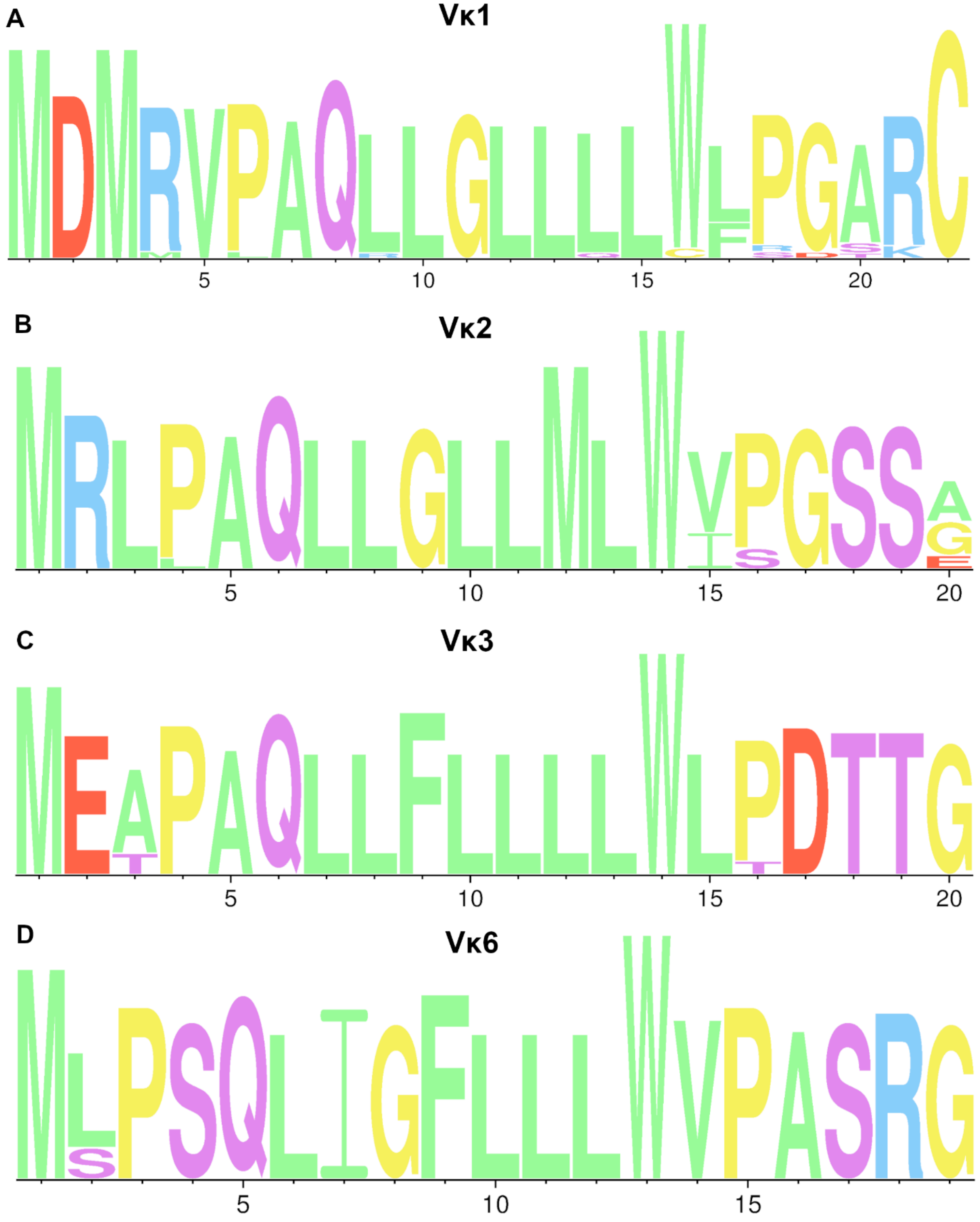

**Supplementary Figure 1.** Consensus sequence analysis of Vκ amino acid sequences using WebLogo (Crooks et al., 2004). Only Vκ1-3 and Vκ6 were shown as both Vκ4 and Vκ5 lacked sufficient sequences for consensus alignment. Blue, Red, Purple, Yellow and Green colors code for the amino acids with positive, negative charges, polar uncharged, special, and hydrophobic side chains, respectively.

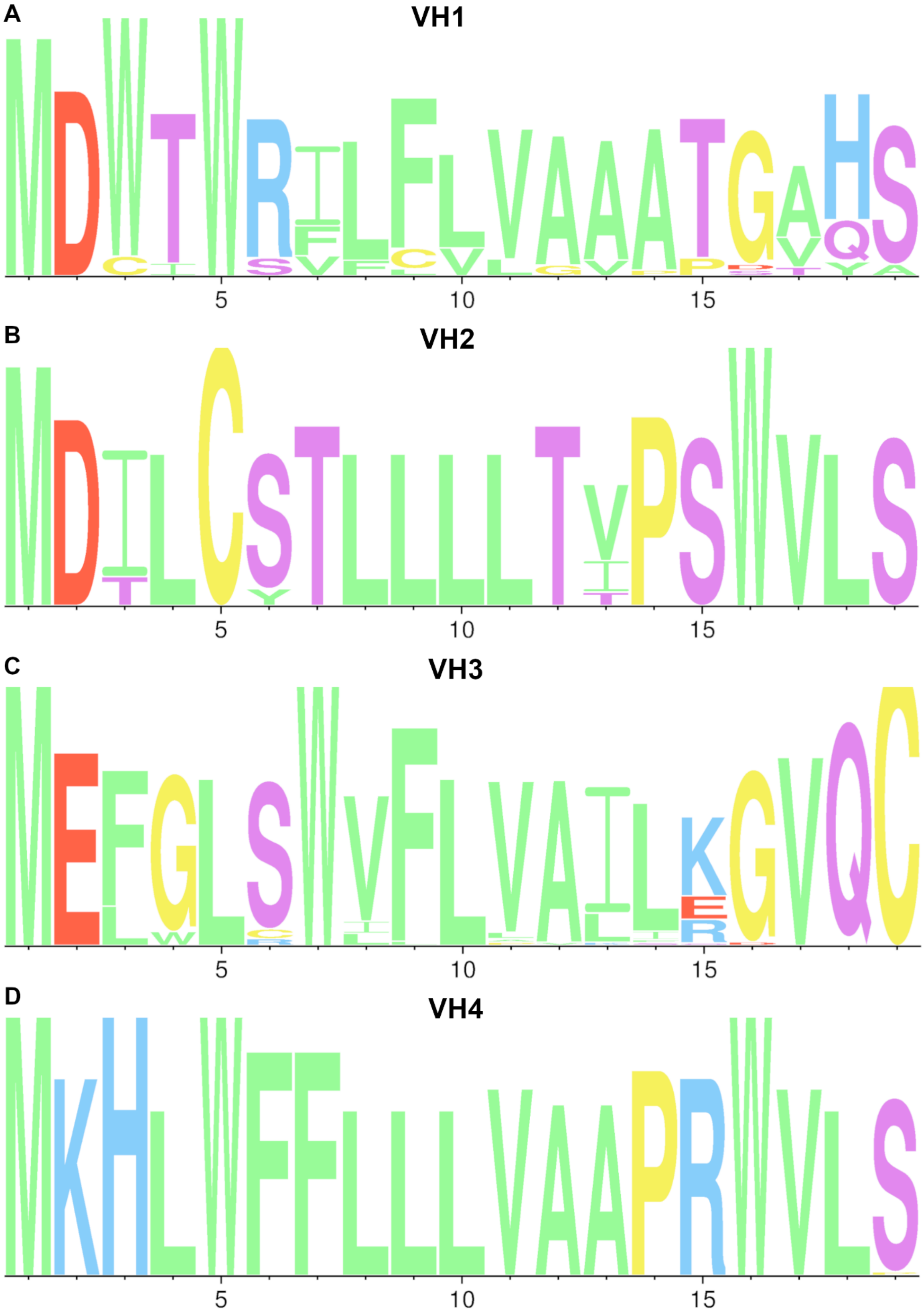
A
VH1
5
10
15
B
VH2
5
10
15
C
VH3
5
10
15
D
VH4
5
10
15

**Supplementary Figure 2.** Consensus sequence analysis of VH amino acid sequences using WebLogo (Crooks et al., 2004). Only VH1-4 were represented here as VH5-7 lacked sufficient sequences for consensus alignment. Blue, Red, Purple, Yellow and Green colors code for the amino acids with positive, negative charges, polar uncharged, special, and hydrophobic side chains, respectively

#### 2 Supplementary Tables

**Supplementary Table 2.** Full amino acid counts of Pertuzumab variants

| Pertuzumab | Amino acid | | | | | | | | | | | | | | | | | | | |
|---|---|---|---|---|---|---|---|---|---|---|---|---|---|---|---|---|---|---|---|---|
| | F | H | I | K | L | M | T | V | W | A | C | D | E | G | N | P | Q | R | S | Y |
| Vκ1\|VH1 | 50 | 24 | 30 | 96 | 110 | 18 | 116 | 130 | 26 | 66 | 32 | 60 | 58 | 84 | 48 | 92 | 68 | 32 | 164 | 66 |
| Vκ2\|VH1 | 48 | 24 | 30 | 94 | 110 | 18 | 108 | 136 | 26 | 64 | 32 | 60 | 62 | 86 | 48 | 96 | 66 | 32 | 162 | 66 |
| Vκ3\|VH1 | 52 | 24 | 28 | 92 | 116 | 14 | 116 | 128 | 26 | 70 | 32 | 56 | 64 | 84 | 48 | 94 | 66 | 34 | 162 | 66 |
| Vκ4\|VH1 | 50 | 24 | 34 | 94 | 106 | 16 | 112 | 136 | 26 | 68 | 32 | 62 | 60 | 84 | 50 | 92 | 68 | 32 | 160 | 66 |
| Vκ5\|VH1 | 56 | 26 | 38 | 94 | 100 | 16 | 116 | 126 | 26 | 72 | 32 | 58 | 64 | 84 | 54 | 94 | 66 | 30 | 154 | 66 |
| Vκ6\|VH1 | 52 | 24 | 32 | 100 | 106 | 14 | 116 | 132 | 26 | 66 | 32 | 60 | 64 | 80 | 50 | 92 | 68 | 30 | 160 | 64 |
| Vκ1\|VH2 | 48 | 22 | 34 | 96 | 124 | 16 | 124 | 124 | 24 | 64 | 32 | 60 | 56 | 80 | 52 | 98 | 64 | 34 | 158 | 64 |
| Vκ2\|VH2 | 46 | 22 | 34 | 94 | 124 | 16 | 116 | 130 | 24 | 62 | 32 | 60 | 60 | 82 | 52 | 102 | 62 | 34 | 156 | 64 |
| Vκ3\|VH2 | 50 | 22 | 32 | 92 | 130 | 12 | 124 | 122 | 24 | 68 | 32 | 56 | 62 | 80 | 52 | 100 | 62 | 36 | 156 | 64 |
| Vκ4\|VH2 | 48 | 22 | 38 | 94 | 120 | 14 | 120 | 130 | 24 | 66 | 32 | 62 | 58 | 80 | 54 | 98 | 64 | 34 | 154 | 64 |
| Vκ5\|VH2 | 54 | 24 | 42 | 94 | 114 | 14 | 124 | 120 | 24 | 70 | 32 | 58 | 62 | 80 | 58 | 100 | 62 | 32 | 148 | 64 |
| Vκ6\|VH2 | 50 | 22 | 36 | 100 | 120 | 12 | 124 | 126 | 24 | 64 | 32 | 60 | 62 | 76 | 54 | 98 | 64 | 32 | 154 | 62 |
| Vκ1\|VH3 | 52 | 22 | 30 | 92 | 120 | 16 | 104 | 130 | 24 | 66 | 32 | 58 | 60 | 90 | 52 | 92 | 64 | 38 | 164 | 66 |
| Vκ2\|VH3 | 50 | 22 | 30 | 90 | 120 | 16 | 96 | 136 | 24 | 64 | 32 | 58 | 64 | 92 | 52 | 96 | 62 | 38 | 162 | 66 |
| Vκ3\|VH3 | 54 | 22 | 28 | 88 | 126 | 12 | 104 | 128 | 24 | 70 | 32 | 54 | 66 | 90 | 52 | 94 | 62 | 40 | 162 | 66 |
| Vκ4\|VH3 | 52 | 22 | 34 | 90 | 116 | 14 | 100 | 136 | 24 | 68 | 32 | 60 | 62 | 90 | 54 | 92 | 64 | 38 | 160 | 66 |
| Vκ5\|VH3 | 58 | 24 | 38 | 90 | 110 | 14 | 104 | 126 | 24 | 72 | 32 | 56 | 66 | 90 | 58 | 94 | 62 | 36 | 154 | 66 |
| Vκ6\|VH3 | 54 | 22 | 32 | 96 | 116 | 12 | 104 | 132 | 24 | 66 | 32 | 58 | 66 | 86 | 54 | 92 | 64 | 36 | 160 | 64 |
| Vκ1\|VH4 | 52 | 24 | 32 | 92 | 120 | 16 | 110 | 130 | 26 | 62 | 32 | 58 | 54 | 84 | 50 | 96 | 70 | 36 | 168 | 64 |
| Vκ2\|VH4 | 50 | 24 | 32 | 90 | 120 | 16 | 102 | 136 | 26 | 60 | 32 | 58 | 58 | 86 | 50 | 100 | 68 | 36 | 166 | 64 |
| Vκ3\|VH4 | 54 | 24 | 30 | 88 | 126 | 12 | 110 | 128 | 26 | 66 | 32 | 54 | 60 | 84 | 50 | 98 | 68 | 38 | 166 | 64 |
| Vκ4\|VH4 | 52 | 24 | 36 | 90 | 116 | 14 | 106 | 136 | 26 | 64 | 32 | 60 | 56 | 84 | 52 | 96 | 70 | 36 | 164 | 64 |
| Vκ5\|VH4 | 58 | 26 | 40 | 90 | 110 | 14 | 110 | 126 | 26 | 68 | 32 | 56 | 60 | 84 | 56 | 98 | 68 | 34 | 158 | 64 |
| Vκ6\|VH4 | 54 | 24 | 34 | 96 | 116 | 12 | 110 | 132 | 26 | 62 | 32 | 58 | 60 | 80 | 52 | 96 | 70 | 34 | 164 | 62 |
| Vκ1\|VH5 | 46 | 24 | 36 | 98 | 116 | 20 | 106 | 124 | 24 | 66 | 32 | 58 | 60 | 86 | 48 | 92 | 64 | 32 | 170 | 66 |
| Vκ2\|VH5 | 44 | 24 | 36 | 96 | 116 | 20 | 98 | 130 | 24 | 64 | 32 | 58 | 64 | 88 | 48 | 96 | 62 | 32 | 168 | 66 |
| Vκ3\|VH5 | 48 | 24 | 34 | 94 | 122 | 16 | 106 | 122 | 24 | 70 | 32 | 54 | 66 | 86 | 48 | 94 | 62 | 34 | 168 | 66 |

| | | | | | | | | | | | | | | | | | | | |
|---|---|---|---|---|---|---|---|---|---|---|---|---|---|---|---|---|---|---|---|
| Vκ4\|VH5 | 46 | 24 | 40 | 96 | 112 | 18 | 102 | 130 | 24 | 68 | 32 | 60 | 62 | 86 | 50 | 92 | 64 | 32 | 166 | 66 |
| Vκ5\|VH5 | 52 | 26 | 44 | 96 | 106 | 18 | 106 | 120 | 24 | 72 | 32 | 56 | 66 | 86 | 54 | 94 | 62 | 30 | 160 | 66 |
| Vκ6\|VH5 | 48 | 24 | 38 | 102 | 112 | 16 | 106 | 126 | 24 | 66 | 32 | 58 | 66 | 82 | 50 | 92 | 64 | 30 | 166 | 64 |
| Vκ1\|VH6 | 52 | 22 | 38 | 90 | 122 | 14 | 110 | 124 | 24 | 60 | 32 | 58 | 56 | 82 | 54 | 98 | 72 | 34 | 168 | 64 |
| Vκ2\|VH6 | 50 | 22 | 38 | 88 | 122 | 14 | 102 | 130 | 24 | 58 | 32 | 58 | 60 | 84 | 54 | 102 | 70 | 34 | 166 | 64 |
| Vκ3\|VH6 | 54 | 22 | 36 | 86 | 128 | 10 | 110 | 122 | 24 | 64 | 32 | 54 | 62 | 82 | 54 | 100 | 70 | 36 | 166 | 64 |
| Vκ4\|VH6 | 52 | 22 | 42 | 88 | 118 | 12 | 106 | 130 | 24 | 62 | 32 | 60 | 58 | 82 | 56 | 98 | 72 | 34 | 164 | 64 |
| Vκ5\|VH6 | 58 | 24 | 46 | 88 | 112 | 12 | 110 | 120 | 24 | 66 | 32 | 56 | 62 | 82 | 60 | 100 | 70 | 32 | 158 | 64 |
| Vκ6\|VH6 | 54 | 22 | 40 | 94 | 118 | 10 | 110 | 126 | 24 | 60 | 32 | 58 | 62 | 78 | 56 | 98 | 72 | 32 | 164 | 62 |
| Vκ1\|VH7 | 52 | 24 | 30 | 90 | 116 | 16 | 110 | 124 | 26 | 68 | 32 | 58 | 60 | 90 | 50 | 92 | 64 | 36 | 166 | 66 |
| Vκ2\|VH7 | 50 | 24 | 30 | 88 | 116 | 16 | 102 | 130 | 26 | 66 | 32 | 58 | 64 | 92 | 50 | 96 | 62 | 36 | 164 | 66 |
| Vκ3\|VH7 | 54 | 24 | 28 | 86 | 122 | 12 | 110 | 122 | 26 | 72 | 32 | 54 | 66 | 90 | 50 | 94 | 62 | 38 | 164 | 66 |
| Vκ4\|VH7 | 52 | 24 | 34 | 88 | 112 | 14 | 106 | 130 | 26 | 70 | 32 | 60 | 62 | 90 | 52 | 92 | 64 | 36 | 162 | 66 |
| Vκ5\|VH7 | 58 | 26 | 38 | 88 | 106 | 14 | 110 | 120 | 26 | 74 | 32 | 56 | 66 | 90 | 56 | 94 | 62 | 34 | 156 | 66 |
| Vκ6\|VH7 | 54 | 24 | 32 | 94 | 112 | 12 | 110 | 126 | 26 | 68 | 32 | 58 | 66 | 86 | 52 | 92 | 64 | 34 | 162 | 64 |
| **MEDAIN** | **52** | **24** | **34** | **92** | **116** | **14** | **110** | **128** | **24** | **66** | **32** | **58** | **62** | **84** | **52** | **96** | **64** | **34** | **162** | **64** |

**Supplementary Table 3.** Full amino acid counts of Pertuzumab variants

| Trastuzumab | Amino acid | | | | | | | | | | | | | | | | | | | |
|---|---|---|---|---|---|---|---|---|---|---|---|---|---|---|---|---|---|---|---|---|
| | F | H | I | K | L | M | T | V | W | A | C | D | E | G | N | P | Q | R | S | Y |
| Vκ1\|VH1 | 46 | 28 | 30 | 96 | 110 | 18 | 120 | 130 | 28 | 70 | 32 | 60 | 58 | 82 | 46 | 92 | 66 | 36 | 162 | 62 |
| Vκ2\|VH1 | 44 | 28 | 30 | 94 | 110 | 18 | 114 | 136 | 28 | 68 | 32 | 60 | 62 | 86 | 46 | 96 | 64 | 34 | 158 | 62 |
| Vκ3\|VH1 | 48 | 28 | 28 | 92 | 116 | 14 | 122 | 128 | 28 | 74 | 32 | 56 | 64 | 84 | 46 | 94 | 64 | 36 | 158 | 62 |
| Vκ4\|VH1 | 46 | 28 | 34 | 94 | 106 | 16 | 118 | 136 | 28 | 72 | 32 | 62 | 60 | 84 | 48 | 92 | 66 | 34 | 156 | 62 |
| Vκ5\|VH1 | 52 | 30 | 38 | 94 | 100 | 16 | 122 | 126 | 28 | 76 | 32 | 58 | 64 | 84 | 52 | 94 | 64 | 32 | 150 | 62 |
| Vκ6\|VH1 | 48 | 28 | 32 | 100 | 106 | 14 | 122 | 132 | 28 | 70 | 32 | 60 | 64 | 80 | 48 | 92 | 66 | 32 | 156 | 60 |
| Vκ1\|VH2 | 44 | 26 | 34 | 96 | 124 | 16 | 128 | 124 | 26 | 68 | 32 | 60 | 56 | 78 | 50 | 98 | 62 | 38 | 156 | 60 |
| Vκ2\|VH2 | 42 | 26 | 34 | 94 | 124 | 16 | 122 | 130 | 26 | 66 | 32 | 60 | 60 | 82 | 50 | 102 | 60 | 36 | 152 | 60 |
| Vκ3\|VH2 | 46 | 26 | 32 | 92 | 130 | 12 | 130 | 122 | 26 | 72 | 32 | 56 | 62 | 80 | 50 | 100 | 60 | 38 | 152 | 60 |
| Vκ4\|VH2 | 44 | 26 | 38 | 94 | 120 | 14 | 126 | 130 | 26 | 70 | 32 | 62 | 58 | 80 | 52 | 98 | 62 | 36 | 150 | 60 |
| Vκ5\|VH2 | 50 | 28 | 42 | 94 | 114 | 14 | 130 | 120 | 26 | 74 | 32 | 58 | 62 | 80 | 56 | 100 | 60 | 34 | 144 | 60 |
| Vκ6\|VH2 | 46 | 26 | 36 | 100 | 120 | 12 | 130 | 126 | 26 | 68 | 32 | 60 | 62 | 76 | 52 | 98 | 62 | 34 | 150 | 58 |
| Vκ1\|VH3 | 48 | 26 | 32 | 92 | 116 | 16 | 110 | 128 | 26 | 74 | 32 | 58 | 60 | 88 | 50 | 92 | 62 | 40 | 162 | 62 |
| Vκ2\|VH3 | 46 | 26 | 32 | 90 | 116 | 16 | 104 | 134 | 26 | 72 | 32 | 58 | 64 | 92 | 50 | 96 | 60 | 38 | 158 | 62 |
| Vκ3\|VH3 | 50 | 26 | 30 | 88 | 122 | 12 | 112 | 126 | 26 | 78 | 32 | 54 | 66 | 90 | 50 | 94 | 60 | 40 | 158 | 62 |
| Vκ4\|VH3 | 48 | 26 | 36 | 90 | 112 | 14 | 108 | 134 | 26 | 76 | 32 | 60 | 62 | 90 | 52 | 92 | 62 | 38 | 156 | 62 |

| | | | | | | | | | | | | | | | | | | | |
|---|---|---|---|---|---|---|---|---|---|---|---|---|---|---|---|---|---|---|---|
| Vκ5\|VH3 | 54 | 28 | 40 | 90 | 106 | 14 | 112 | 124 | 26 | 80 | 32 | 56 | 66 | 90 | 56 | 94 | 60 | 36 | 150 | 62 |
| Vκ6\|VH3 | 50 | 26 | 34 | 96 | 112 | 12 | 112 | 130 | 26 | 74 | 32 | 58 | 66 | 86 | 52 | 92 | 62 | 36 | 156 | 60 |
| Vκ1\|VH4 | 48 | 28 | 32 | 92 | 120 | 16 | 114 | 130 | 28 | 66 | 32 | 58 | 54 | 82 | 48 | 96 | 68 | 40 | 166 | 60 |
| Vκ2\|VH4 | 46 | 28 | 32 | 90 | 120 | 16 | 108 | 136 | 28 | 64 | 32 | 58 | 58 | 86 | 48 | 100 | 66 | 38 | 162 | 60 |
| Vκ3\|VH4 | 50 | 28 | 30 | 88 | 126 | 12 | 116 | 128 | 28 | 70 | 32 | 54 | 60 | 84 | 48 | 98 | 66 | 40 | 162 | 60 |
| Vκ4\|VH4 | 48 | 28 | 36 | 90 | 116 | 14 | 112 | 136 | 28 | 68 | 32 | 60 | 56 | 84 | 50 | 96 | 68 | 38 | 160 | 60 |
| Vκ5\|VH4 | 54 | 30 | 40 | 90 | 110 | 14 | 116 | 126 | 28 | 72 | 32 | 56 | 60 | 84 | 54 | 98 | 66 | 36 | 154 | 60 |
| Vκ6\|VH4 | 50 | 28 | 34 | 96 | 116 | 12 | 116 | 132 | 28 | 66 | 32 | 58 | 60 | 80 | 50 | 96 | 68 | 36 | 160 | 58 |
| Vκ1\|VH5 | 42 | 28 | 36 | 98 | 116 | 20 | 110 | 124 | 26 | 70 | 32 | 58 | 60 | 84 | 46 | 92 | 62 | 36 | 168 | 62 |
| Vκ2\|VH5 | 40 | 28 | 36 | 96 | 116 | 20 | 104 | 130 | 26 | 68 | 32 | 58 | 64 | 88 | 46 | 96 | 60 | 34 | 164 | 62 |
| Vκ3\|VH5 | 44 | 28 | 34 | 94 | 122 | 16 | 112 | 122 | 26 | 74 | 32 | 54 | 66 | 86 | 46 | 94 | 60 | 36 | 164 | 62 |
| Vκ4\|VH5 | 42 | 28 | 40 | 96 | 112 | 18 | 108 | 130 | 26 | 72 | 32 | 60 | 62 | 86 | 48 | 92 | 62 | 34 | 162 | 62 |
| Vκ5\|VH5 | 48 | 30 | 44 | 96 | 106 | 18 | 112 | 120 | 26 | 76 | 32 | 56 | 66 | 86 | 52 | 94 | 60 | 32 | 156 | 62 |
| Vκ6\|VH5 | 44 | 28 | 38 | 102 | 112 | 16 | 112 | 126 | 26 | 70 | 32 | 58 | 66 | 82 | 48 | 92 | 62 | 32 | 162 | 60 |
| Vκ1\|VH6 | 48 | 26 | 38 | 90 | 122 | 14 | 114 | 124 | 26 | 64 | 32 | 58 | 56 | 80 | 52 | 98 | 70 | 38 | 166 | 60 |
| Vκ2\|VH6 | 46 | 26 | 38 | 88 | 122 | 14 | 108 | 130 | 26 | 62 | 32 | 58 | 60 | 84 | 52 | 102 | 68 | 36 | 162 | 60 |
| Vκ3\|VH6 | 50 | 26 | 36 | 86 | 128 | 10 | 116 | 122 | 26 | 68 | 32 | 54 | 62 | 82 | 52 | 100 | 68 | 38 | 162 | 60 |
| Vκ4\|VH6 | 48 | 26 | 42 | 88 | 118 | 12 | 112 | 130 | 26 | 66 | 32 | 60 | 58 | 82 | 54 | 98 | 70 | 36 | 160 | 60 |
| Vκ5\|VH6 | 54 | 28 | 46 | 88 | 112 | 12 | 116 | 120 | 26 | 70 | 32 | 56 | 62 | 82 | 58 | 100 | 68 | 34 | 154 | 60 |
| Vκ6\|VH6 | 50 | 26 | 40 | 94 | 118 | 10 | 116 | 126 | 26 | 64 | 32 | 58 | 62 | 78 | 54 | 98 | 70 | 34 | 160 | 58 |
| Vκ1\|VH7 | 48 | 28 | 30 | 90 | 116 | 16 | 114 | 124 | 28 | 72 | 32 | 58 | 60 | 88 | 48 | 92 | 62 | 40 | 164 | 62 |
| Vκ2\|VH7 | 46 | 28 | 30 | 88 | 116 | 16 | 108 | 130 | 28 | 70 | 32 | 58 | 64 | 92 | 48 | 96 | 60 | 38 | 160 | 62 |
| Vκ3\|VH7 | 50 | 28 | 28 | 86 | 122 | 12 | 116 | 122 | 28 | 76 | 32 | 54 | 66 | 90 | 48 | 94 | 60 | 40 | 160 | 62 |
| Vκ4\|VH7 | 48 | 28 | 34 | 88 | 112 | 14 | 112 | 130 | 28 | 74 | 32 | 60 | 62 | 90 | 50 | 92 | 62 | 38 | 158 | 62 |
| Vκ5\|VH7 | 54 | 30 | 38 | 88 | 106 | 14 | 116 | 120 | 28 | 78 | 32 | 56 | 66 | 90 | 54 | 94 | 60 | 36 | 152 | 62 |
| Vκ6\|VH7 | 50 | 28 | 32 | 94 | 112 | 12 | 116 | 126 | 28 | 72 | 32 | 58 | 66 | 86 | 50 | 92 | 62 | 36 | 158 | 60 |
| **MEDAIN** | **48** | **28** | **34** | **92** | **116** | **14** | **114** | **128** | **26** | **70** | **32** | **58** | **62** | **84** | **50** | **96** | **62** | **36** | **158** | **60** |
